# A new T Cell Receptor in Squamata Reptiles

**DOI:** 10.1101/2023.09.04.556186

**Authors:** Francisco Gambón-Deza

## Abstract

Squamata exhibit a loss of genes for the gamma/delta T-lymphocyte receptor chains and a significant decrease in the number of V genes at the TRBV locus. Through genome analysis, I have discovered a new locus that contains V, J, C, and TM genes that have a similar structure to the classical TCR chains. This gene is viable, as demonstrated by the presence of messenger RNAs in the transcriptomes. Analyses using the AlphaFold2 program indicate that the deduced protein chain is associated with the alpha chain of the TCR. I have named this new chain “epsilon,” and it forms a new TCR alpha/epsilon. Evolutionarily, the epsilon chain arose from a duplication of the beta chain gene at the time of the divergence of amphibians and reptiles and has since been specifically maintained in Squamata.

## 1. Introduction

The process of antigen recognition by classic T-lymphocyte receptor is complex (Garcia et al., 1999). Native antigens are not recognized by this receptor; they must first be processed within the cell and presented by major histocompatibility complex (MHC) proteins (Bjorkman et al., 1987; Birnbaum et al., 2014). Dual recognition occurs through the interaction of the MHC molecule and the antigen with the T-lymphocyte receptor (Garcia et al., 1996). T-lymphocyte receptor are composed by two chains called alpha and beta. A second type of receptor was later discovered composed of other two chains named gamma and delta. Each one of them is expressed in different T lymphocytes defining two populations. T cells with the alpha-beta receptor is the most prevalent and has the ability to recognize molecules in MHC, while T cells with gamma-delta receptors are not MHC-restricted in recognition (Brenner et al., 1987; Raulet, 1989).

The genes that code for these receptors share a similar structure. Each chain is composed of a gene for the constant chain, a set of short-sequence genes (D and J) to induce variability, and a cluster of V genes encoding the variable sections. Additionally, there are one or two small cytoplasmic exons and an exon for the transmembrane region (Davis, 1990).

These receptors first became visible in jawed vertebrates. These genes were created at the same time as the genes for antibodies and MHC and shape the specific acquired immunity system. In every species, the receptor with the alpha-beta chains has persisted. It’s interesting to note that the gamma-delta receptor is missing in several fish species and is absent from the Squamata evolutionary branch (Flajnik, 2002; Rast & Litman, 1994; Olivieri et al., 2014).

In reptiles, there are two main evolutionary lines, the Squamata and the Archosaurs. The former are geckos, lizards, iguanas, and snakes, while the latter are turtles, crocodiles, and birds. Within the immune system, significant variations are detected in the Squamata. They do not have gamma-delta T cells (Olivieri et al., 2014; Gambón-Deza & Olivieri, 2018; Morrissey et al., 2022). Within the genes encoding antibody chains, all have genes for lambda-light chains. Kappa chain genes are no longer present in snakes but are found in Squamata taxa with more ancient origins, such as geckos (Gambón-Deza et al., 2012). At the level of heavy chain genes, five isotypes of three classes of immunoglobulins have been found (1 IgM, 1 IgD, and 2 or 3 IgY isotypes)(Olivieri et al., 2021). Another feature is the low number of V regions in the TCR beta chain locus and the large number of genes for MHC antigens in serpents (Olivieri et al., 2020).

In a survey of squamata genomes, a new locus characteristic of translocon genes (V/J/C genes) I found. These loci code for an amino acid chain similar to the TCR chains. This gene may help to explain the gene changes found in these species.

## 2. Material and Methods

The entire study is based on freely available data. The sequences of the genomes are in the NCBI and on the web at https://vgp.github.io/. We use our applications developed in Python to obtain the sequences of interest. For the general study of the sequences, the Biopython library was used (Cock et al., 2009). The exons of the constant regions of the TCR chains genes were performed with a machine learning (tensorflow (Abadi, 2016)) trained for their recognition. Other publications express the specific operation (Mirete-Bachiller et al., 2021). The exons for the variable regions were obtained with the Vgenextractor program (Olivieri et al., 2021). These applications extract the possible target exon. The amino acid sequences are analyzed by machine learning, and the target sequences are obtained.

The messenger RNA sequences were obtained from the RNAseq files deposited in the NCBI. The sequences were aligned to the reference genome on the web https://usegalaxy.org/ whith program hisat2 (Kim et al., 2019) and visualized with the program IGV (Thorvaldsdóttir et al., 2013).

The 3D representations were generated with the alphafold2 software (Jumper et al., 2021). We used a version on the cloud with the notebook available at https://colab.research.google.com/github/sokrypton/ColabFold/blob/main/AlphaFold2.ipynb#scrollTo=_sztQyz29DIC.

The graphic representation of the exons was also done with a simple python application using the genomediagram library (Pritchard et al., 2006).

The amino acid sequences obtained were aligned with the MAFFT program (Katoh et al., 2005). The trees were built with the Fasttree program (Price et al., 2010) using the LG matrix (Le & Gascuel, 2008) and gamma parameter. The tree visualisation was done with the Figtree program (http://tree.bio.ed.ac.uk/software/figtree/)

## 3. Results

Squamate reptiles, which include lizards and snakes, have experienced the loss of genes for major immune system molecules compared to Archosauromorpha reptiles. Some species of squamates have lost the kappa light chain immunoglobulin, and all species have lost the genes for the gamma and delta chains of the second type of T-lymphocyte receptor (Olivieri et al., 2014). This raises questions about how squamates compensate for the absence of these immune system components and how this evolutionary process has occurred. Although there is no definitive answer yet, researchers have found that large deletions in the genome of squamate reptiles, such as the sleepy lizard, have resulted in the removal of genes necessary to produce γ/δ T cells, which are an important part of the immune system in most vertebrates (Morrissey et al., 2022). However, the genes encoding the α/β T cell receptor (TCR) chains in squamates do not appear to have increased in complexity to compensate for the loss of γ/δ T cells. Further research is needed to understand the compensatory mechanisms or variations that may have evolved in squamate reptiles to maintain their immune system functionality despite the loss of these genes.

### 3.1. Genes V

The Vgeneextractor program is trained with sequences from IGHV, IGKV, IGLV, TR(A/D)V, and TRGV genes. Each exon is assigned a class by the computer based on the percentage of accuracy it receives. If there are unknown sequences similar to some of the classes, they can be assigned to one of these classes with a low level of confidence. One surprising finding is the localization of some genes assigned to TRAV and TRBV. Some of these genes are clustered at a locus located on a different chromosome from where the constant region gene is located. All these sequences on other chromosomes are grouped into a cluster. These sequences indeed correspond to V region sequences. As this V genes are similar to the V genes of T cell receptors, we refer to them as V epsilon of the TREV locus. However, the variability between them is low, so the decision was made to retrain the Vgeneextractor program by adding this class (V*ϵ*)

### 3.2 A new T cell receptor chain

In all genomes from all Squamata species this V*ϵ* regions are detected, but not in Archosauro-mopha (turtles, crocodiles and birds). Between 2 and 22 V regions are found per species. All of them very similar in sequence. As can be seen in the table 1 no TRGV genes and very few TRBV genes are detected in any species (Vgeneextractor does not detect 100% of V genes, which may explain why some species do not obtain V genes in the TRBV region (Olivieri et al., 2013)). In previous publications it was already mentioned that no genes for the TRγ/δ receptor are found (Olivieri et al., 2014; Gambón-Deza & Olivieri, 2018). This has already been confirmed by others (Morrissey et al., 2022).

**Table 1:** Number of genes Vs.

|  | IGHV | IGKV | IGLV | TRA/DV | TRBV | TRGV | TREV |
| --- | --- | --- | --- | --- | --- | --- | --- |
| *Eublepharis macularius* | 180 | 3 | 123 | 23 | 2 | - | 1 |
| *Paroedura picta* | 48 | - | 49 | 8 | 6 | - | - |
| *Gekko japonicus* | 184 | 2 | 115 | 23 | 10 | - | - |
| *Phrynocephalus versicolor* | 44 | 11 | 32 | 18 | 3 | - | 1 |
| *Pogona vitticeps* | 86 | 12 | 52 | 12 | 2 | - | 5 |
| *Anolis carolinensis* | 105 | 23 | 37 | 19 | 8 | - | 4 |
| *Varanus salvator* | 52 | - | 75 | 1 | 1 | - | 0 |
| *Varanus komodoensis* | 218 | - | 151 | 65 | 4 | - | 22 |
| COLUBRIDAE |  |  |  |  |  |  |  |
| *Arizona elegans* | 227 | - | 112 | 17 | 1 | - | 4 |
| *Thermophis baileyi* | 159 | - | 73 | 12 | 3 | - | 5 |
| *Pytas mucosa* | 215 | - | 72 | 19 | - | - | 5 |
| *Diadophis punctatus* | 211 | - | 68 | 21 | 2 | - | 5 |
| *Pituophis catenifer* | 149 | - | 53 | 12 | 1 | - | 2 |
| *Thamnophis elegans* | 286 | - | 299 | 21 | 1 | - | 7 |
| VIPERIDAE |  |  |  |  |  |  |  |
| *Bothrops jararaca* | 246 | - | 66 | 28 | 3 | - | 6 |
| *Crotalus viridis* | 70 | - | 44 | 11 | 2 | - | 6 |
| *Crotalus adamanteus* | 146 | - | 57 | 19 | 2 | - | 6 |
| *Vipera latastei* | 104 | - | 45 | 15 | 3 | - | 5 |
| *Protobothrops mucrosquamatus* | 138 | - | 54 | 17 | 4 | - | 10 |
| ELAPIDAE |  |  |  |  |  |  |  |
| *Emydocephalus ijimae* | 49 | - | 35 | 7 | - | - | 5 |
| *Notechis scutatus* | 87 | - | 26 | 13 | - | - | 2 |
| *Hidrophis curtus* | 87 | - | 98 | 21 | - | - | 7 |
| *Hidrophis cyanocinctus* | 82 | - | 45 | 12 | 1 | - | 5 |
| *Naja naja* | 99 | - | 77 | 14 | - | - | 14 |
| *Bungarus multicinctus* | 144 | - | 148 | 38 | - | - | 9 |
| PYTHONIDAE |  |  |  |  |  |  |  |
| *Python molurus* | 81 | - | 45 | 28 | 3 | - | 7 |
| BOIDAE |  |  |  |  |  |  |  |
| *Charina bottae* | 103 | - | 33 | 15 | 2 | - | 2 |

It was decided to look for the constant region that accompanies these Vs. To do so, we studied the presence of these Vs in messenger RNAs that should carry the constant region. A search was carried out for the presence of messenger RNAs in the databases available at NCBI. It is found complete sequences from the TSA database (for example GIKH01051440.1, GIMS01001238.1). In this way it was possible to obtain sequences of the constant region and extend the study of the genome of these species. In Figure 1, the results obtained in the snake *Bungarus multicinctus* are presented. From the transcriptome SRR11177805 were 144 sequences compatible. These long sequences (sequencing with paciphic bioscience) were aligned to the region of the chromosomal contig CM042838.1. The presence of six V regions, one exon J, one exon for the constant region, another exon for the transmembrane region, and a cytoplasmic exon with the 3’ untranslated region were detected. This structure is very similar to that of the T-lymphocyte receptor genes.

**Figure 1.**
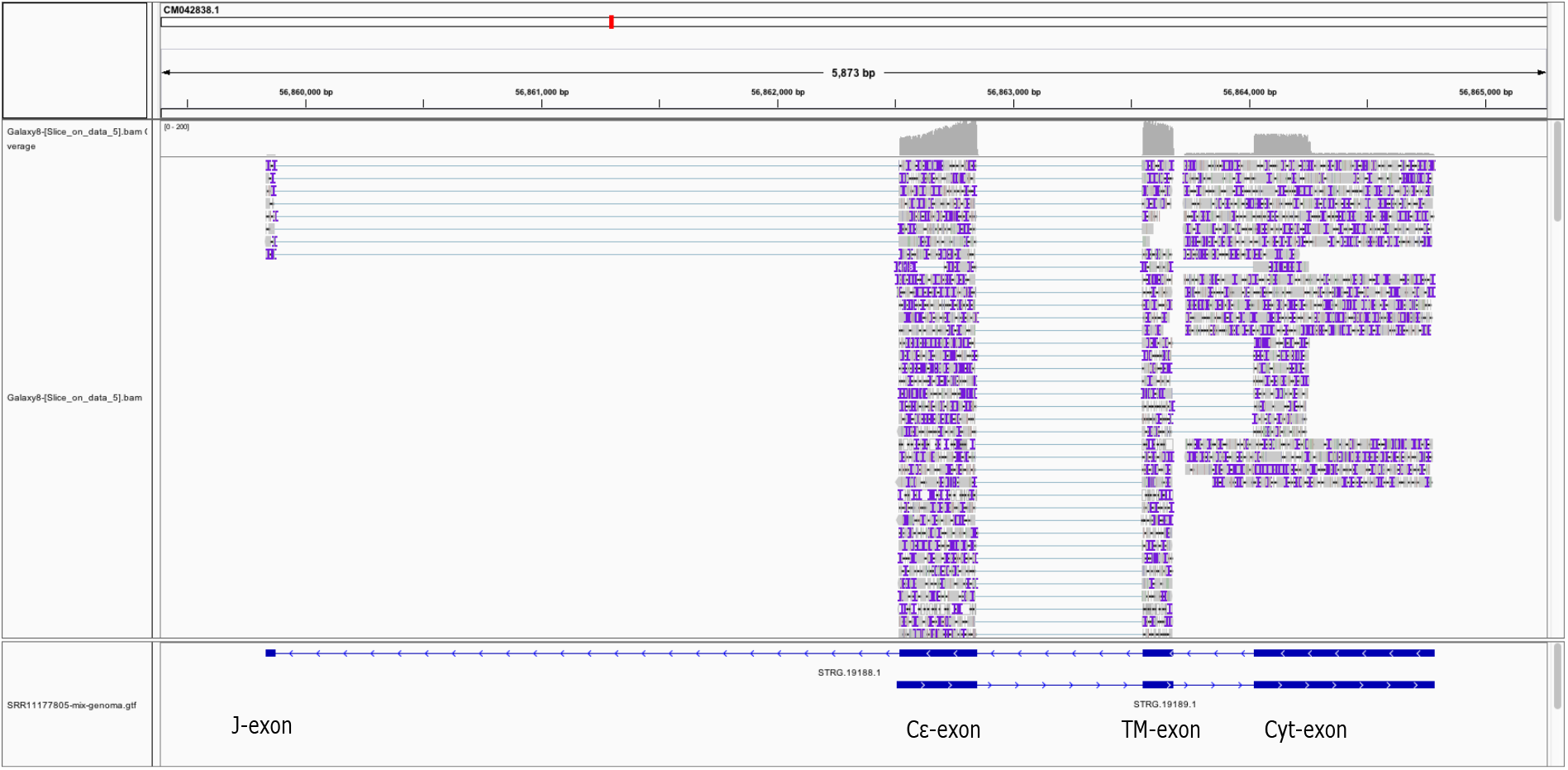
Results obtained in the alignment of long reads obtained with the Paciphic Biscience platform of the snake species *Bungarus multicinctus* (SRR11177805). The sequences were aligned with the minimap2 program and visualised with the IGV program. A view of the region where the constant region is located is presented, showing the presence of J minigenes, one exon for the C region, one for the transmembrane region and one for the cytoplasmic region.

By using Blast we were able to find sequences of the constant region and determined that just like the Vϵ regions it is present in Squamata and not in other reptiles (Figure 4). The constant region is highly conserved between species. There are 5 cysteines in the constant region that form disulphide bridges deduced from the three-dimensional structure explained below.

We try to determine its evolutionary origin. In all Squamata studied present it. They are not detected in archosaurs or amphibians. In a phylogenetic tree with light chains and other types of T cell receptor constant chain sequences, the TCR epsilon chain constant region appears to have a closer origin to the TCR beta chain constant region. Its origin goes back to the time of the divergence of reptiles with amphibians, and it was only kept in the evolutionary line of the Squamata (figure 2).

**Figure 2.**
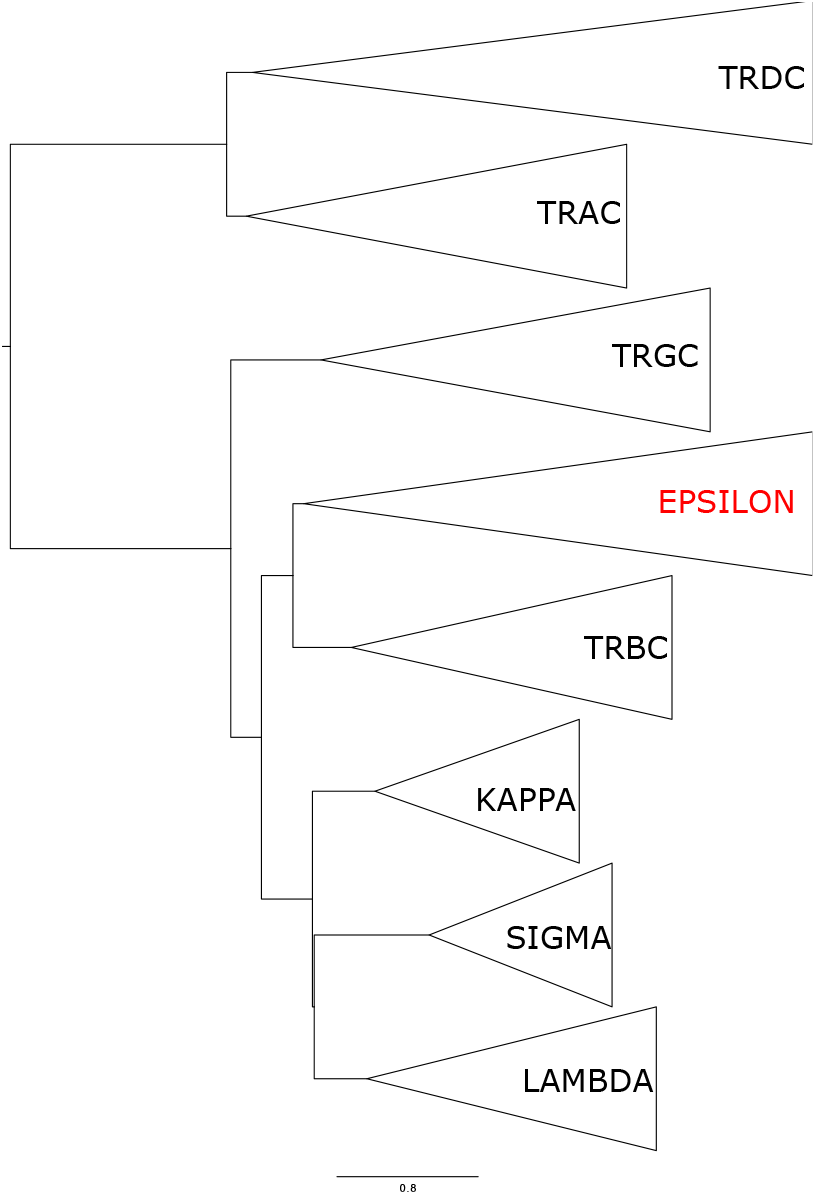
360 sequences of the constant regions of reptiles and amphibians of light chains (Kappa, Lambda, and Sigma) and constant regions of the TCR chains (TRAC, TRBC, TRGC and TREC-Epsilon) were aligned and the phylogenetic tree was made with the Fasttree program (LG matrix, gamma paramiter). The tree suggests an origin origin of the trbc trbc.

**Figure 3.**
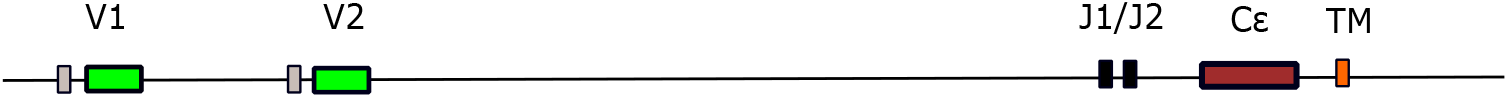
Genomic structure of the loci coding for the Epsilon chain of the TCR found in the snake *Notechis scutatus*. Sequence ID: UWZA01000165.1, Length: 1253545. The loci is located in chain minus between 800000 and 850000 nt

**Figure 4.**
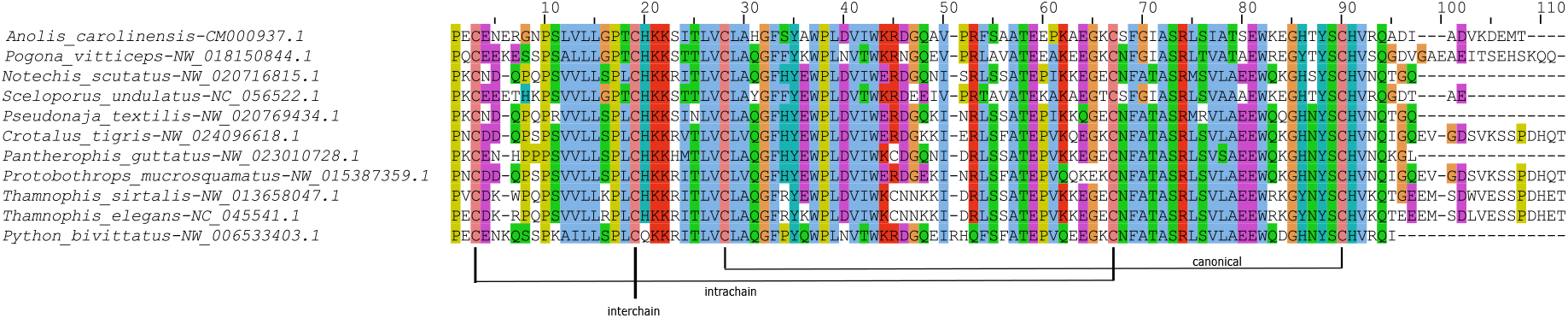
Alignment of deduced sequences of the gene coding for the constant region of the epsilon chain

### 3.3 3D structure

To find out if this chain is associated with one of the alpha or beta chains of the T lymphocyte receptor or if it forms homodimers, we use the three-dimensional structure deductions with the alphafold2 program (Figure 5. When we ask if this chain can form a homodimer, a non-suggestive three-dimensional structure emerges since the sequences tend to pair in reverse directions or there are no clear interaction sites between the chains. The epsilon chain forms a viable configuration with the alpha chain, existing in proximity and forming an inter chain bridge in the constant domain and leaving the V regions in a position similar to that described in the TCR with the CDR3 regions of each chain close to each other. With the beta chain, it forms a non-viable structure with distant constant domains without inter-chain covalent bonding.

**Figure 5.**
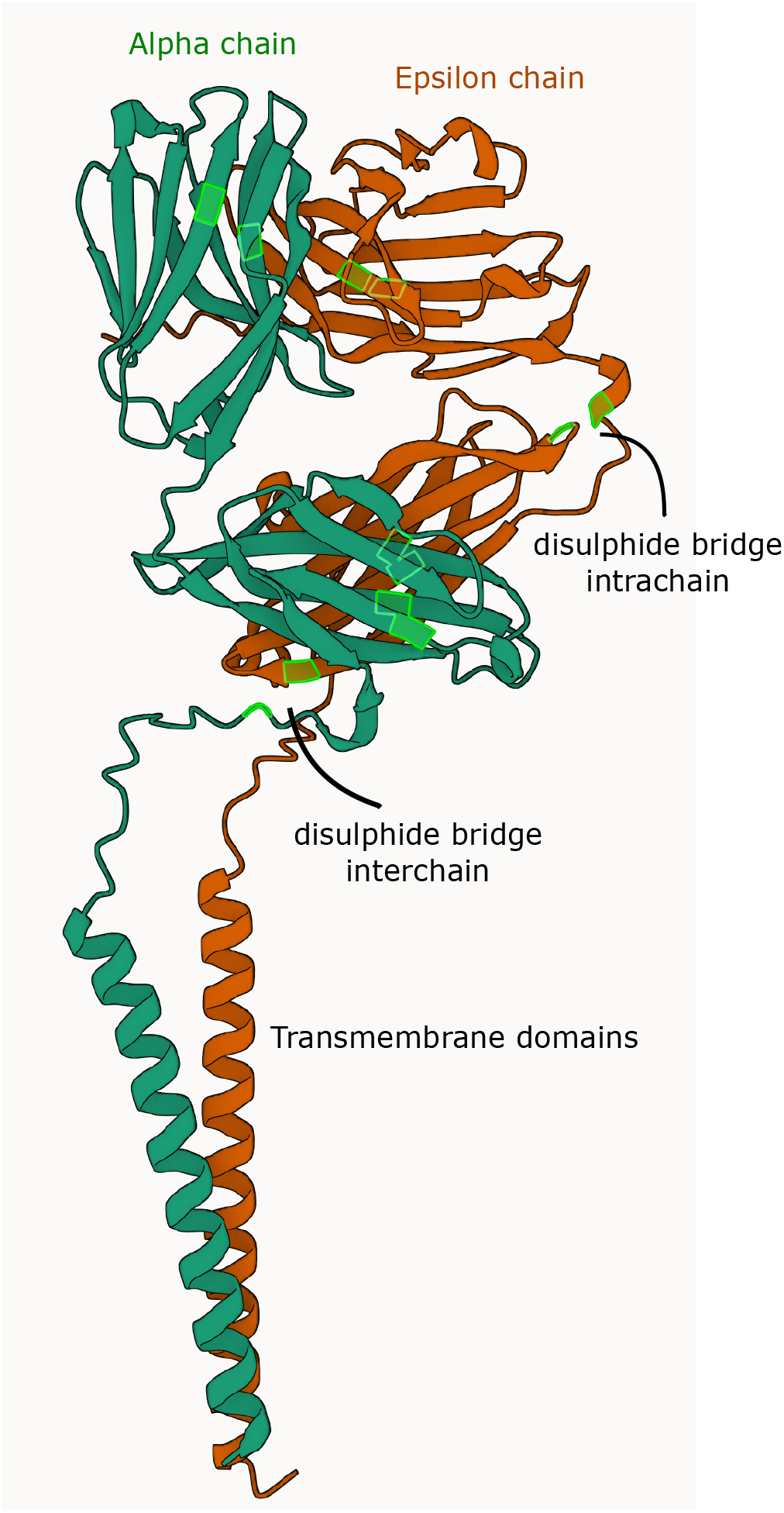
3D structure of the alpha/epsilon hetero-dimer deduced with the Alphafold program. The sequences correspond to those obtained from the transcriptomes of *Bungarus multicinctus*. Deduced disulphide bridges are indicated.

## 4. Discussion

The evolutionary lineage of reptilian Squamata has unique features in the major genes responsible for immunity. The immunoglobulin genes contain genes for heavy and light chains, but many species do not have a gene for the kappa chain. Genes for the alpha and beta chains of the TCR are present, but no Squamata species have genes for the gamma and delta chains. In addition, a significant decrease in genes at the TRBV locus is detected.

In this study, we describe the presence of a new gene that may help us understand the radical changes that have occurred. The gene is very similar to a T-lymphocyte receptor chain. The alphafold program indicates that this chain is highly suggestive of conforming to heterodimers with the alpha chain of the TCR. Thus, we have a new TCR made up of alpha and epsilon chains. The epsilon chain has its origin in a duplication of the beta chain, which supports the results. It may also explain the decline in the number of genes at the TRBV locus, probably due to compensation in the generation of new loci.

Antigen recognition is expected to be the same. The association of CDRs in the three-dimensional structure suggests this. The new receptor must correspond to a significant population of T lymphocytes as it is widely expressed in the tissues studied.

Could it be responsible for the loss of the gamma delta complex? Bearing in mind that the presence of this new receptor correlates inversely with the presence of the gamma/delta receptor, the data highly suggest that it is responsible. Additionally, it has similarities with the gamma/delta TCR as the latter expresses Vδ sequences that are found within the chromosomal segment where Vα is located.

It may also explain the low number of Vβ regions present in Squamata. This would indicate that there is usurpation of functions and that this chain with the alpha chain has the functions of both alpha-beta and gamma-delta.

The results suggest evolutionary dynamics. A duplication of the T-lymphocyte receptor beta chain locus must have occurred. For unknown reasons, it ended up on another chromosome. This chain led to the formation of an alpha-epsilon receptor. The functionality of this new receptor gave them an evolutionary advantage, and the gamma-delta receptor was no longer needed and was eliminated from the genome. It is more difficult to infer whether this also conditioned the decrease in V genes in the germ line of the TRB locus as there are other significant variations in the genomes of these species, notably the increase in genes for the MHC class I and class II beta-chain antigens (Olivieri et al., 2020).

## References

Abadi, M. (2016). Tensorflow: learning functions at scale. In Proceedings of the 21st ACM SIGPLAN International Conference on Functional Programming (pp. 1–1).

Birnbaum, M. E., Mendoza, J. L., Sethi, D. K., Dong, S., Glanville, J., Dobbins, J., Ö zkan, E., Davis, M. M., Wucherpfennig, K. W., & Garcia, K. C. (2014). Deconstructing the peptide-mhc specificity of t cell recognition. Cell, 157, 1073–1087.

Bjorkman, P., Saper, M., Samraoui, B., Bennett, W., Strominger, J., & Wiley, D. (1987). The foreign antigen binding site and t cell recognition regions of class i histocompatibility antigens. Nature, 329, 512–518.

Brenner, M. B., McLean, J., Scheft, H., Riberdy, J., Ang, S.-L., Seidman, J., Devlin, P., & Krangel, M. S. (1987). Two forms of the t-cell receptor γ protein found on peripheral blood cytotoxic t lymphocytes. Nature, 325, 689–694.

Cock, P. J., Antao, T., Chang, J. T., Chapman, B. A., Cox, C. J., Dalke, A., Friedberg, I., Hamelryck, T., Kauff, F., Wilczynski, B. et al. (2009). Biopython: freely available python tools for computational molecular biology and bioinformatics. Bioinformatics, 25, 1422–1423.

Davis, M. M. (1990). T cell receptor gene diversity and selection. Annual review of biochemistry, 59, 475–496.

Flajnik, M. F. (2002). Comparative analyses of immunoglobulin genes: surprises and portents. Nature Reviews Immunology, 2, 688–698.

Gambón-Deza, F., & Olivieri, D. N. (2018). Immunoglobulin and t cell receptor genes in chinese crocodile lizard shinisaurus crocodilurus. Molecular immunology, 101, 160–166.

Gambón-Deza, F., Sánchez-Espinel, C., Mirete-Bachiller, S., & Magadán-Mompó, S. (2012). Snakes antibodies. Developmental & Comparative Immunology, 38, 1–9.

Garcia, K. C., Degano, M., Stanfield, R. L., Brunmark, A., Jackson, M. R., Peterson, P. A., Teyton, L., & Wilson, I. A. (1996). An αβ t cell receptor structure at 2.5 å and its orientation in the tcr-mhc complex. Science, 274, 209–219.

Garcia, K. C., Teyton, L., & Wilson, I. A. (1999). Structural basis of t cell recognition. Annual review of immunology, 17, 369.

Jumper, J., Evans, R., Pritzel, A., Green, T., Figurnov, M., Ronneberger, O., Tunyasuvunakool, K., Bates, R., Žídek, A., Potapenko, A. et al. (2021). Highly accurate protein structure prediction with alphafold. Nature, 596, 583–589.

Katoh, K., Kuma, K.-i., Toh, H., & Miyata, T. (2005). Mafft version 5: improvement in accuracy of multiple sequence alignment. Nucleic acids research, 33, 511–518.

Kim, D., Paggi, J. M., Park, C., Bennett, C., & Salzberg, S. L. (2019). Graph-based genome alignment and genotyping with hisat2 and hisat-genotype. Nature biotechnology, 37, 907–915.

Le, S. Q., & Gascuel, O. (2008). An improved general amino acid replacement matrix. Molecular biology and evolution, 25, 1307–1320.

Mirete-Bachiller, S., Olivieri, D. N., & Gambón-Deza, F. (2021). Gouania willdenowi is a teleost fish without immunoglobulin genes. Molecular Immunology, 132, 102–107.

Morrissey, K. A., Sampson, J. M., Rivera, M., Bu, L., Hansen, V. L., Gemmell, N. J., Gardner, M. G., Bertozzi, T., & Miller, R. D. (2022). Comparison of reptilian genomes reveals deletions associated with the natural loss of γδ t cells in squamates. The Journal of Immunology, 208, 1960–1967.

Olivieri, D., Faro, J., von Haeften, B., Sánchez-Espinel, C., & Gambón-Deza, F. (2013). An automated algorithm for extracting functional immunologic v-genes from genomes in jawed vertebrates. Immunogenetics, 65, 691–702.

Olivieri, D., Mirete-Bachiller, S., & Gambón-Deza, F. (2020). Mhc class i and ii genes in serpentes. bioRxiv, (pp. 2020–06).

Olivieri, D., Von Haeften, B., Sánchez-Espinel, C., Faro, J., & Gambón-Deza, F. (2014). Genomic v exons from whole genome shotgun data in reptiles. Immunogenetics, 66, 479–492.

Olivieri, D. N., Mirete-Bachiller, S., & Gambón-Deza, F. (2021). Insights into the evolution of ig genes in amphibians and reptiles. Developmental & Comparative Immunology, 114, 103868.

Price, M. N., Dehal, P. S., & Arkin, A. P. (2010). Fasttree 2–approximately maximum-likelihood trees for large alignments. PloS one, 5, e9490.

Pritchard, L., White, J. A., Birch, P. R., & Toth, I. K. (2006). Genomediagram: a python package for the visualization of large-scale genomic data. Bioinformatics, 22, 616–617.

Rast, J. P., & Litman, G. W. (1994). T-cell receptor gene homologs are present in the most primitive jawed vertebrates. Proceedings of the National Academy of Sciences, 91, 9248–9252.

Raulet, D. H. (1989). The structure, function, and molecular genetics of the gamma/delta t cell receptor. Annual review of immunology, 7, 175–207.

Thorvaldsdóttir, H., Robinson, J. T., & Mesirov, J. P. (2013). Integrative genomics viewer (igv): high-performance genomics data visualization and exploration. Briefings in bioinformatics, 14, 178–192.

